# Perm1 Gene Therapy Mitigates PRDM16-Associated Cardiomyopathy

**DOI:** 10.64898/2026.04.13.718330

**Authors:** Omid MT Rouzbehani, Sophie Stephens, Ben Werbner, Marta W. Szulik, Bo Sun, Mira Hua, Shinya Watanabe, Ava Leonelli, Michael Goodman, Ryan Bia, Crystal F. Davey, Martin G Golkowski, Sarah Franklin, Andrew P. Landstrom, Sihem Boudina

## Abstract

**Background:** Pathogenic variants in PR domain containing 16 (*PRDM16*) cause pediatric and adult cardiomyopathies characterized by ventricular dilation, systolic dysfunction, and impaired metabolic maturation. Cardiac deficiency of PRDM16 alters metabolic gene expression and long-chain fatty acid (FA) metabolites. However, the downstream mediators involved are not well characterized. Furthermore, whether improving mitochondrial FA metabolism can prevent PRDM16-associated cardiomyopathy is currently unknown.

**Methods:** *In vivo* and *in vitro* approaches using patient-induced pluripotent stem cell-derived cardiomyocytes (iPSC-CMs) and mouse models with *Prdm16* deletion/mutation were employed. Transcriptomics and proteomics analyses were conducted, and adeno-associated virus (AAV)-mediated therapy was tested.

**Results:** Here, we show that a defect in FA metabolism is an early hallmark of PRDM16 cardiac deficiency. We show, for the first time, that PERM1 is a direct downstream target of PRDM16 and is involved in the regulation of FA metabolism through coordinated action with PGC1α. Most importantly, neonatal delivery of AAV9–*Perm1* in cardiac-specific *Prdm16* knockout (*Prdm16* cKO) mice markedly improved contractile parameters, reduced left ventricular (LV) dilation, and extended survival. These cardioprotective effects of PERM1 gene therapy occurred independent of restoring FA oxidation. Transcriptional and proteomic analyses of AAV–*Perm1*-treated *Prdm16* cKO mice demonstrated significant improvements in mitochondrial cristae architecture, preservation of sarcomere organization, reduced cardiomyocyte apoptosis, attenuated myocardial fibrosis, and diminished cardiac remodeling.

**Conclusions:** We identify PERM1 as a direct downstream effector of PRDM16 and uncover a previously unrecognized PRDM16–PGC1α–PERM1 axis essential for FA metabolic regulation in the heart. *Perm1* gene therapy ameliorated PRDM16-associated cardiomyopathy through post-transcriptional mechanisms involving preservation of mitochondrial and sarcomere integrity. The current study provides preclinical evidence suggesting that *Perm1* gene therapy may be a promising therapeutic target to improve the cardiac outcomes of patients affected by pathogenic *PRDM16* variants.

## Introduction

Cardiomyopathies comprise a diverse group of myocardial disorders that impair cardiac structure and function and are major contributors to heart failure (HF), arrhythmias, and sudden cardiac death^1^. Although many forms are acquired, a substantial proportion, particularly in children, are caused by pathogenic genetic variants^2^. These inherited cardiomyopathies, most transmitted in an autosomal dominant pattern, give rise to distinct phenotypes, including dilated, hypertrophic, restrictive, and LV non-compaction (LVNC) cardiomyopathies^3^. Pediatric genetic cardiomyopathies remain a leading indication for heart transplantation, with nearly 40% of symptomatic patients progressing to end-stage HF within two years of diagnosis, yet many individuals still lack a definitive molecular diagnosis^4^. This underscores an unmet need to identify disease-causing genes and mechanistic pathways that may yield new therapeutic opportunities.

PRDM16, a zinc-finger transcription factor with lysine methyltransferase activity, has been associated with LVNC and dilated cardiomyopathy (DCM) phenotypes in patients with chromosome 1p36 deletion syndrome^5^. Our group further demonstrated that PRDM16 deletion in 1p36 deletion syndrome patients is causal for these cardiac phenotypes^6^. In mice, several knockouts of *Prdm16* have been generated using different *Cre* drivers. Using *Myh6-Cre*, we and others showed that cardiac-specific deletion of *Prdm16* caused DCM in mice^6,7^. Cibi et al^8^ used *Mesp-Cre* to delete *Prdm16* in cardiac progenitors and showed that the KO mice develop progressive heart failure, characterized by increased cardiac hypertrophy, reduced fatty acid gene expression and metabolite levels, and impaired mitochondrial energetics when challenged with a high-fat diet. Using two cardiomyocyte-specific *Cre* drivers (*Xmlc2-Cre* and *cTnT-Cre)*, Wu et al^9^showed that *Prdm16* deletion resulted in LVNC, DCM, and perinatal mortality and mechanistically linked these phenotypes to altered chamber specifications. Smooth muscle cell deletion of *Prdm16* using *Sm22a-Cre* caused early onset cardiomyopathy in mice^10^. Our group developed a knock-in *Prdm16 Prdm16^Q187X/WT^* mouse that exhibits embryonic lethality associated with severe LVNC, a phenotype also seen in a patient carrying the same mutation^11^. Finally, the Klassen group^12^ used a monoallelic, global *Prdm16* mutant mouse (*Prdm16^csp1/wt^*) and showed that these animals exhibited reduced cardiac metabolites, including those involved in the tricarboxylic acid (TCA) cycle, glycerol, amino acids, and pentose phosphate pathway. All these germline deletions and mutations in *Prdm16* in mice underscore the crucial role of this transcription factor during embryonic development of the heart.

A common finding in the above-mentioned studies is the consistent alteration in cardiac metabolic pathways following loss of PRDM16. However, the molecular drivers involved in the regulation of cardiac metabolism, downstream of PRDM16 in the heart, are not fully characterized. Furthermore, the contribution of altered cardiac metabolism to the DCM phenotype of *Prdm16* cKO mice is currently unknown. Finally, whether restoring metabolic function may constitute a valid therapeutic approach to attenuate PRDM16-associated cardiomyopathy has never been previously explored.

In this study, we identified PERM1 and PGC1α as downstream transcriptional targets of PRDM16 in the heart, demonstrating that regulation of FA oxidation by PRDM16 requires PGC1α and PERM1. We further show that AAV9-mediated *Perm1* overexpression markedly improved systolic function, limited LV dilation, reduced myocardial cell death, attenuated fibrosis, and enhanced survival in *Prdm16* cKO mice. These protective effects of *Perm1* gene therapy occurred independent of restoring FA oxidation. Rather, *Perm1* gene therapy and improved mitochondrial and sarcomere integrity in the heart. These findings define a mechanistic link between PRDM16 and PGC1α/PERM1 in the regulation of cardiac FA metabolism and establish *Perm1* gene therapy as a protective intervention for PRDM16-associated cardiomyopathy in mice.

## Methods

For detailed information on mouse experimental procedures, refer to Supplemental Material. Primary data, where useful to the general scientific community, are uploaded as supplemental files. The corresponding author can be contacted for any data or reagents used with the appropriate regulatory request documents.

### Experimental Animals

All animal experiments were approved by the University of Utah Institutional Animal Care and Use Committee and conducted in accordance with NIH guidelines. Mice were housed under a 12-hour light–dark cycle with ad libitum access to food and water. Both male and female mice were included. Investigators were blinded to genotype and treatment during data acquisition and analysis unless otherwise specified. Cardiomyocyte-specific *Prdm16* cKO mice were generated by crossing *Prdm16^flox^*^/flox^ mice with *αMHC-Cre* mice, resulting in deletion of exons 6–7 in cardiomyocytes. Littermate *Prdm16^flox^*^/flox^ mice lacking Cre served as wild-type controls. The *Prdm16^Q187X^*^/WT^ line was generated on a C57BL/6NJ background using CRISPR-Cas9, as previously described^13^. Founder mice were backcrossed ≥7 generations, and correct targeting was confirmed by Sanger sequencing with no detectable off-target effects. Male and female mice were included; breeders and animals with procedural complications were excluded. No randomization was performed, as comparisons were genotype- or treatment-defined.

### Statistical Analysis

Data are presented as mean ± SD. Statistical analyses were performed using GraphPad Prism 10.1. Two-group comparisons used two-tailed unpaired *t*-tests. Multiple groups were analyzed by two-way ANOVA followed by Tukey post hoc test. Survival curves were analyzed using log-rank tests. The *p*-value *p* < 0.05 was considered significant. Investigators were blinded to treatment and genotype.

## Results

### Cardiomyocyte deletion of *Prdm16* alters FA metabolism and oxidative phosphorylation gene expression in mice

Cardiomyocyte-specific deletion of *Prdm16* driven by the *αMHC-Cre* transgene resulted in progressive DCM, characterized by reduced ejection fraction (EF), fractional shortening (FS), and cardiac output, and increased LV dimensions in both sexes (**Figure 1A**). To identify early transcriptional changes preceding cardiac dysfunction, RNA-Seq was performed on 8-week-old *Prdm16* cKO and WT hearts. Differential gene expression and pathway analyses revealed significant downregulation of metabolic genes, with the strongest enrichment for oxidative phosphorylation and fatty acid (FA) metabolism (**Figures 1B and 1C, Figures S1A-S1G**), indicating that loss of *Prdm16* disrupts mitochondrial and FA metabolic gene networks prior to the development of contractile dysfunction.

**Figure 1.**
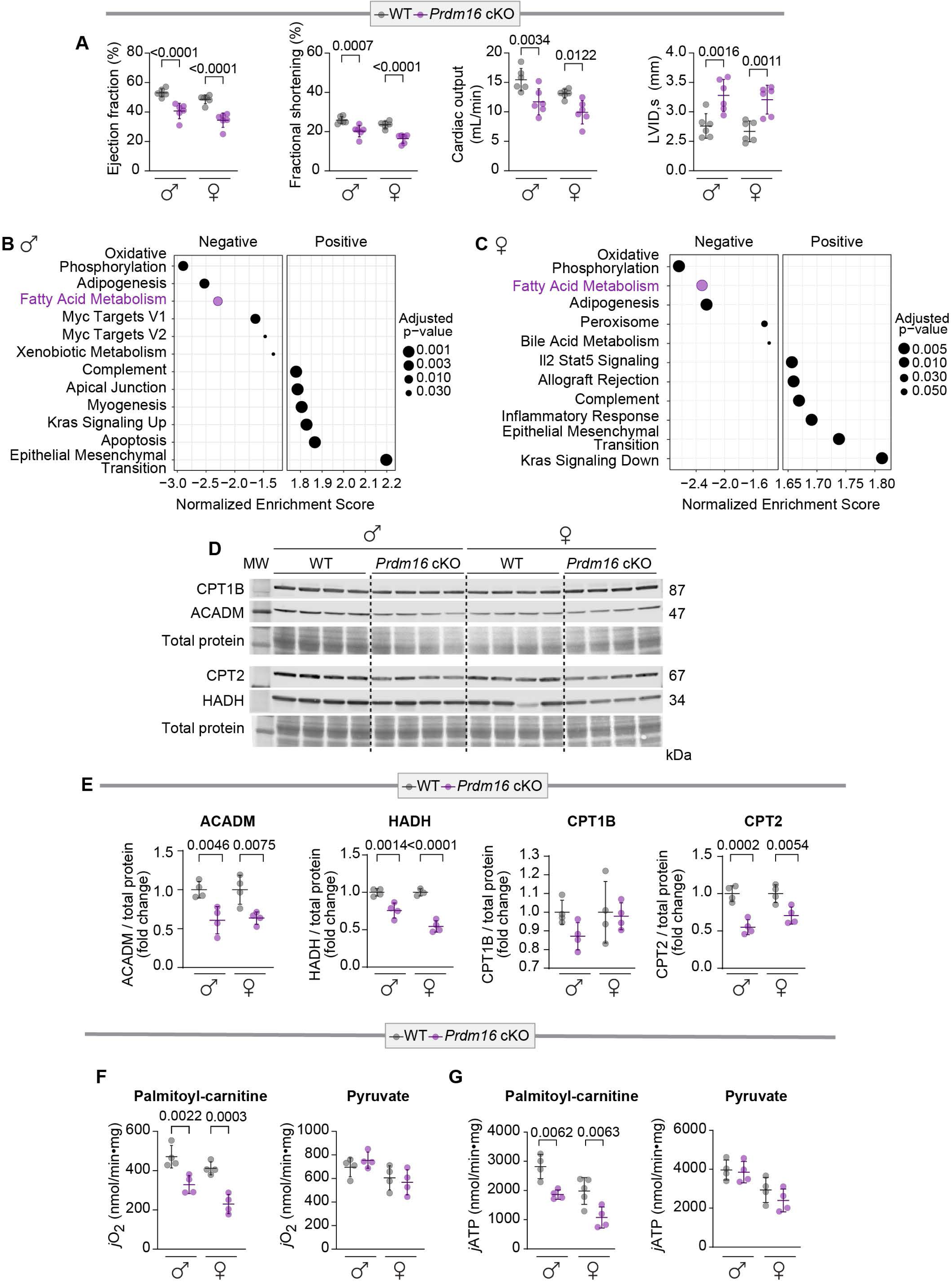
*Prdm16* cKO mice develop dilated cardiomyopathy with impaired fatty acid oxidation and mitochondrial respiration. **(A)** Echocardiographic assessment of 20-week-old male and female WT and *Prdm16* cKO mice demonstrates systolic dysfunction, reduced cardiac output, and increased ventricular dilation in KO animals (N = 6 per group). **(B–C)** Ingenuity pathway analyses in male **(B)** and female **(C)** *Prdm16* cKO hearts reveal significant downregulation of fatty acid (FA) metabolism and oxidative phosphorylation (N = 3 per group). **(D-E)** Western blot analysis of whole heart from 20-week-old male and female mice shows a significant decrease in protein expression of key FA oxidation enzymes, including HADH, ACADM, and CPT2 (N = 4 per group). **(F)** Mitochondrial respiration measured in isolated mitochondria from whole hearts shows significantly reduced oxygen consumption rate (jO₂) with palmitoyl-carnitine (PLC) as substrate, while no significant differences are observed with pyruvate. (**G**) Mitochondrial ATP production rates (jATP) showing a significant reduction in *Prdm16* cKO mice when PLC is used (N = 4 per group). All graphs are mean ± SD; *p*-values are shown on the graph and are calculated by two-way ANOVA followed by a Tukey’s post hoc test.

### Cardiac *Prdm16* deficiency impairs FA oxidation

To test whether the changes in FA metabolic gene expression in *Prdm16* cKO hearts translate into functional impairments in mitochondrial FA utilization, we assessed protein expression of key enzymes of FA transport and oxidation in both male and female hearts at 20 weeks. Western blot analysis showed a significant reduction in Hydroxyacyl-CoA dehydrogenase (HADH), Carnitine palmitoyl transferase 2 (CPT2), and Medium-chain acyl-CoA dehydrogenase (ACADM) compared to WT controls (**Figures 1D and 1E)**. We next measured mitochondrial respiration using pyruvate or palmitoyl-carnitine (PLC). PLC-supported oxygen consumption (JO2) was significantly reduced in *Prdm16* cKO hearts at 20 weeks (**Figure 1F**). In contrast, there was no difference in pyruvate-supported respiration (**Figure 1F**), indicating that *Prdm16* cKO mice have a specific defect in FA utilization. These data are substantiated by the significant reduction in mitochondrial ATP generation when PLC but not pyruvate is used (**Figure 1G**).

### *Perm1* is a direct transcriptional target of PRDM16 in the heart

Since we observed decreased expression of mitochondrial oxidative and FAO genes in *Prdm16* cKO hearts, we first examined mRNA expression of key transcriptional regulators of these pathways. Thus, mRNA expression of *Pparα*, *Ppargc1a* (encoding PGC1α), *Pparγ*, *Esrra* (encoding ERRα), and *Perm1* was measured in *Prdm16* cKO and WT mice. As depicted in **Figures 2A and 2B**, significant downregulation of *Perm1* and *Ppargc1a* mRNA was observed in post-natal day 1 (P1) *Prdm16* cKO hearts and neonatal rat ventricular myocytes (NRVMs) lacking Prdm16. Consistently*, PERM1* and *PPARGC1A* mRNA expression was significantly decreased in human iPSC-CMs carrying the *PRDM16*^Q187X/WT^ mutation compared to the corrected (*PRDM16^cWT/WT^*) isogenic control line (**Figures 2C and 2D**). While PGC1α is a widely studied metabolic coactivator of FAO genes, PERM1 is enriched in striated muscle and has recently been implicated in regulating mitochondrial energetics in the heart. In addition, sustained cardiac PGC1α overexpression has previously been associated with adverse cardiac remodeling^14-17^. Based on its muscle-restricted expression and its recent cardioprotective role, we focused subsequent studies on PERM1. *Perm1* mRNA expression was reduced in *Prdm16*^Q187X/WT^ knock-in mice both at post-natal day 3 and at three months of age **(Figure 2E**). Similarly, PERM1 protein expression was diminished in *Prdm16* cKO hearts from P1 mice (**Figure 2F**). These data demonstrate that PRDM16 deficiency reduces PERM1 expression in murine cardiac tissue and human cells.

**Figure 2.**
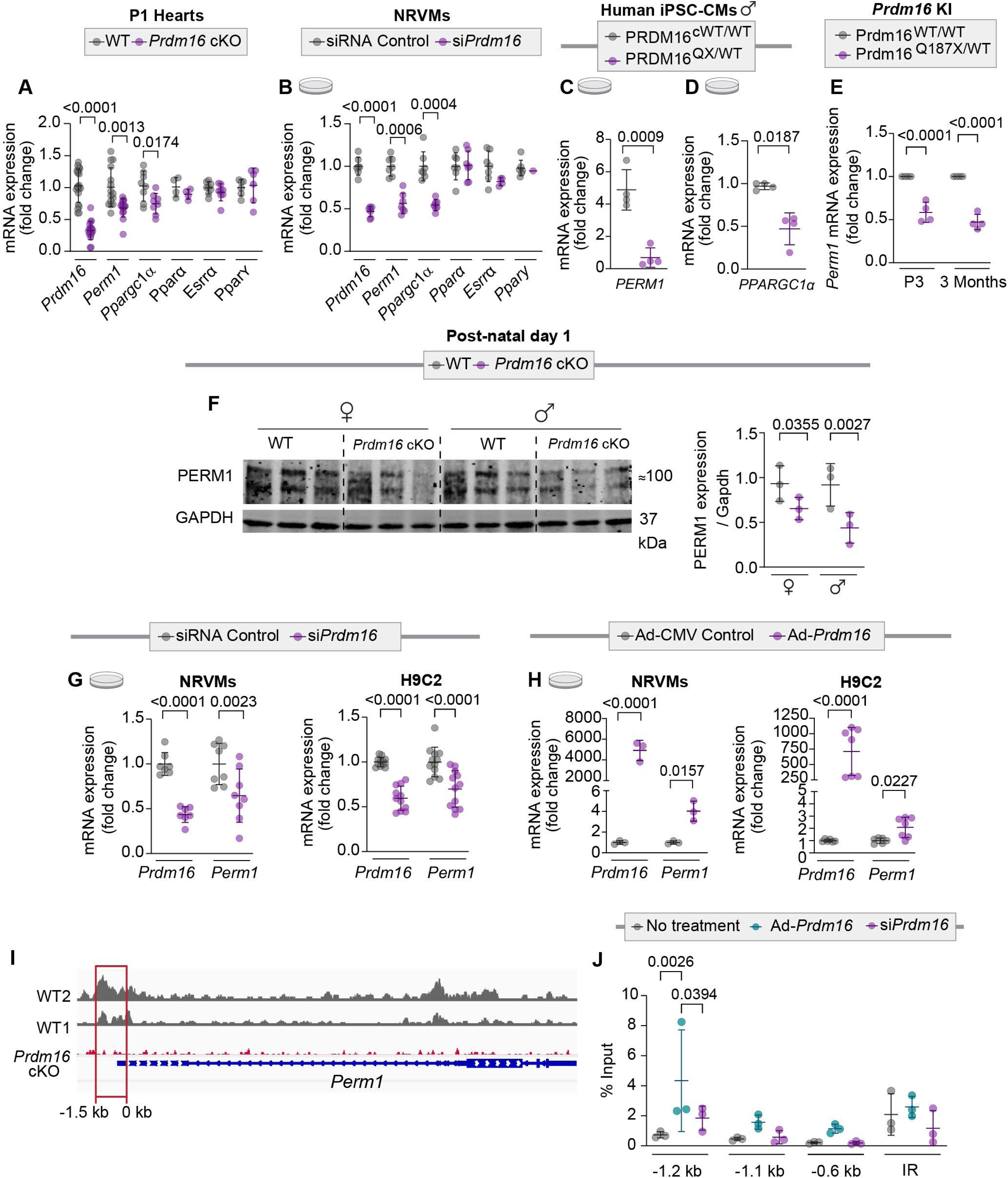
PRDM16 regulates *Perm1* expression in a cell-autonomous manner. (**A**) mRNA expression of *Prdm16*, *Perm1*, and other metabolic regulators (*Ppargc1a*, *Ppara*, *Esrra*, *Pparg*) in postnatal day 1 (P1) hearts from WT and *Prdm16* cKO mice demonstrates a significant reduction in *Perm1* and *Ppargc1a mRNA* expression in KO hearts, with no changes in other metabolic genes (N = 6 per group). (**B**) siRNA-mediated knockdown of *Prdm16* in neonatal rat ventricular myocytes (NRVMs) results in a significant decrease in *Perm1* and *Ppargc1a* mRNA, with minimal effects on other metabolic regulators (N = 6–8 per group). (**C–D**) Human iPSC-derived cardiomyocytes, harboring the *PRDM16* Q187X variant (QX/WT) exhibit significantly reduced *PERM1* (**C**) and *PPARGC1A* (**D**) mRNA expression compared to isogenic corrected controls (cWT/WT) (N = 3–4 per group). (**E**) In a *Prdm16* ^Q187X^ knock-in (KI) mouse model, *Perm1* mRNA expression is significantly reduced at both postnatal day 3 (P3) and 3 months of age (N = 4 per group). (**F)** Western blot analysis of P1 cardiac tissue shows reduced Perm1 protein expression in *Prdm16* cKO hearts relative to WT (N = 3–4 per group). (**G**) Knockdown of *Prdm16* using siRNA in NRVMs and H9c2 cells significantly reduces *Perm1* mRNA expression (N = 6–12 per group). (**H**) Adenoviral overexpression of *Prdm16* (Ad–*Prdm16*) increases *Perm1* mRNA expression in both NRVMs and H9C2 cells compared to Ad–CMV control (N = 6–12 per group). (**I**) Analysis of publicly available ChIP-Seq data (GSE179393) from embryonic day E13.5 hearts shows enrichment of Prdm16 binding at the *Perm1* locus in WT samples, which is absent in *Prdm16* cKO hearts. (**J**) ChIP-qPCR targeting regions within the *Perm1* regulatory locus demonstrates increased Prdm16 occupancy upon overexpression and reduced enrichment following knockdown (N = 3 per group). IR stands for intergenic region. Data are mean ± SD; *p*-values are shown in the panels. Statistical significance is determined using two-way ANOVA followed by Tukey’s post hoc test.

To test whether PRDM16 regulates PERM1 expression in a cell-autonomous manner, we performed loss- and gain-of-function studies using siRNA-mediated *Prdm16* knockdown and adenoviral-mediated *Prdm16* overexpression in H9c2 cells and NRVMs. *Prdm16* siRNA significantly reduced *Perm1* mRNA expression, whereas adenoviral overexpression of *Prdm16* increased *Perm1* expression in both cell types (**Figures 2G and 2H**). To determine how *Prdm16* regulates *Perm1*, we examined publicly available *Prdm16* ChIP-seq data obtained from embryonic hearts at day E13.5 (Accession No. GSE179393)^9^. *Prdm16* shows clear enrichment peaks at the *Perm1* promoter region in WT hearts, which are absent in Prdm16-deficient embryonic hearts (**Figure 2I**). ChIP-qPCR in H9c2 cells further confirmed Prdm16 binding to the *Perm1* promoter, with significant enrichment at −1.2 kb and no enrichment at a random intergenic region (**Figure 2J**). Collectively, these data establish *Perm1* as a downstream transcriptional target of Prdm16 in cardiac cells.

### *Perm1* gene delivery improves cardiac function and reduces LV dilation in *Prdm16* cKO mice

Given that Prdm16 regulates *Perm1* expression in the mouse heart and that *Perm1* was previously shown to regulate both mitochondrial energetics^18-20^ and FAO in the heart ^21^, we hypothesized that *Perm1* overexpression may mitigate cardiomyopathy in *Prdm16* cKO mice through improvement of mitochondrial function and FA metabolism. Indeed, two recent studies from two different laboratories showed that *Perm1* overexpression, prior to pressure overload in mice, improved contractile and mitochondrial function^22,23^. WT and *Prdm16* cKO mice were injected at P1 with a single dose (10¹² viral genomes) of AAV9–*Perm1* or AAV9–GFP, driven by the CMV promoter, through the temporal facial vein **(Figure 3A**). Robust GFP expression was observed in the heart at 20 weeks post-injection, confirming efficient AAV delivery. GFP expression was restricted to the heart, with minimal signal detected in liver, lungs, skeletal muscle and brown adipose tissue (BAT), respectively **(Figure 3B, Figure S2A)**. Consistent with this, AAV9–*Perm1* produced sustained and cardiac-specific Perm1 overexpression, with elevated mRNA and protein levels persisting to 20 weeks post-injection in both female and male hearts (**Figures 3C and 3D, Figures S2B-S2E**). Echocardiography analysis demonstrated that Perm1 overexpression significantly improved cardiac function in female *Prdm16* cKO mice, with higher EF and FS and reduced LV dilation compared with GFP-treated *Prdm16* cKO female mice, reaching values similar to WT controls (**Figures 3E and 3F**). Note that Perm1 overexpression did not affect contractile parameters or cardiac dimensions in female WT mice. Surprisingly, male WT mice receiving AAV9–*Perm1* exhibited a significant 5-10% reduction in EF and FS when compared to AAV9-GFP-treated male WT mice (**Figure S2F)**. This was associated with a significant elevation in heart rate only in male WT mice receiving AAV9–*Perm1* (**Figure S2F**). Despite this adverse effect of AAV9–*Perm1* in WT male mice, the protective effect of overexpressing Perm1 in *Prdm16* cKO male mice was still evident, as shown by a trend toward an increase in EF and FS and reduced LV dilation when compared to AAV9–GFP *Prdm16* cKO male mice (**Figure S2F**).

**Figure 3.**
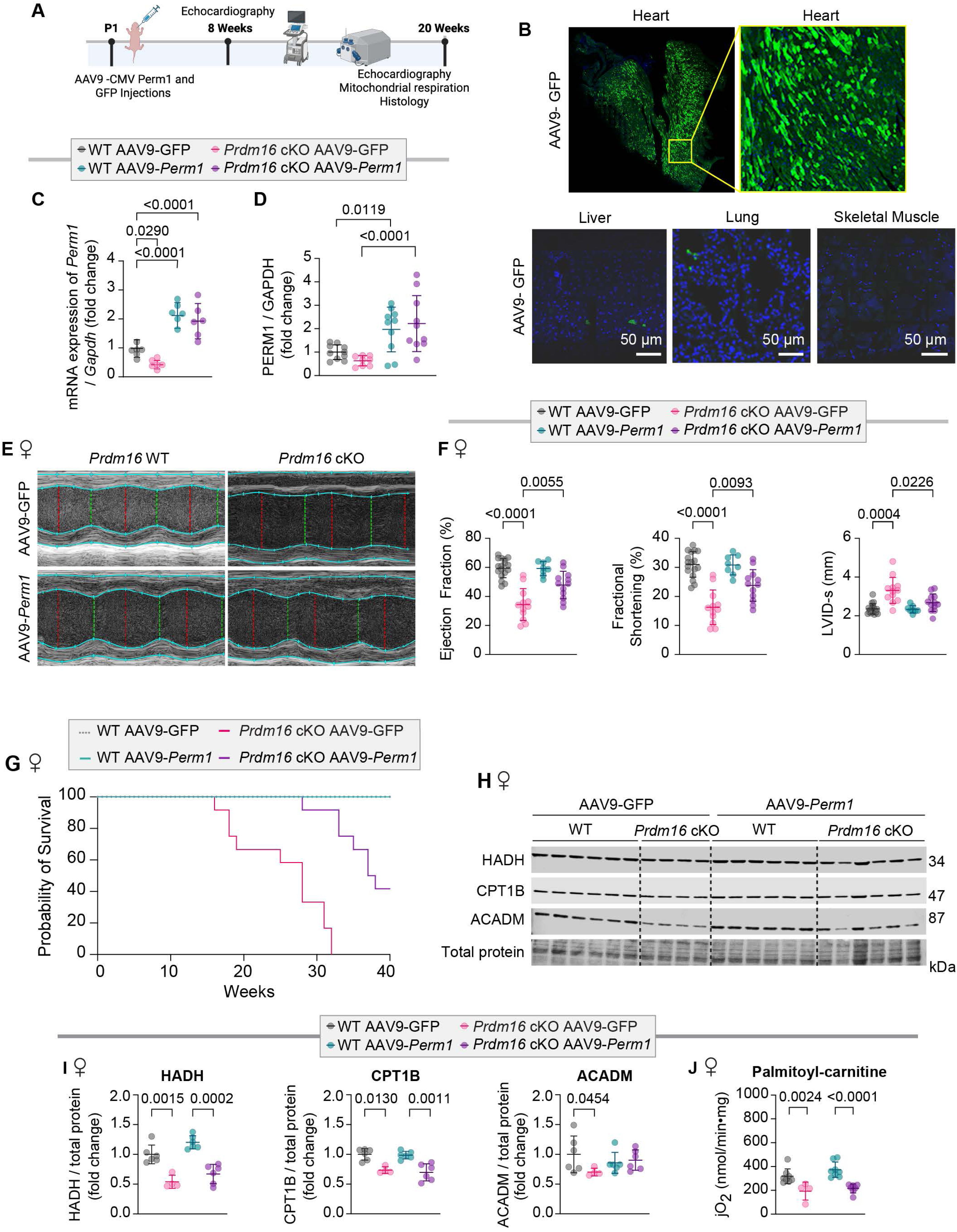
AAV9-mediated *Perm1* gene delivery improves cardiac function and extends survival but does not restore cardiac FA oxidation in *Prdm16* cKO mice. (**A**) Schematic of experimental design. Post neonatal day 1 (P1) WT and *Prdm16* cKO mice received AAV9–CMV–GFP or AAV9–CMV–*Perm1* injections. (**B**) Representative immunofluorescence images demonstrating robust cardiac transduction of AAV9–GFP, with minimal signal detected in liver, lung, and skeletal muscle (scale bars, 50 µm). (**C**) Significant upregulation of *Perm1* mRNA expression in cardiac tissue at 20 weeks post-injection in AAV9–*Perm1*-treated mice compared to AAV9–GFP controls (N = 4–5 per group). (**D**) Cardiac Perm1 protein expression following AAV9–*Perm1* delivery (N = 8-10 per group). (**E**) Representative M-mode echocardiographic images from WT and *Prdm16* cKO mice treated with AAV9–GFP or AAV9–*Perm1*. (**F**) Echocardiographic quantification shows that Perm1 overexpression significantly improves systolic function, as indicated by increased EF and FS, and reduced LVIDs in *Prdm16* cKO mice (N ≥ 8 per group). (**G**) Kaplan–Meier survival analysis demonstrates improved survival in *Prdm16* cKO mice receiving AAV9–*Perm1* (N = 12 per group). (**H**) Western blot analysis of whole heart shows reduced protein expression of FA oxidation (FAO) enzymes (HADH, CPT1B, ACADM) in *Prdm16* cKO mice. (**I**) Quantification of FAO-related proteins normalized to total protein shows minimal changes in AAV9–*Perm1*-treated *Prdm16* cKO mice (N = 3–4 per group). (**J**) Mitochondrial respiration assays using palmitoyl-carnitine (PLC) demonstrate no difference in jO₂ in AAV9-*Perm1*-treated *Prdm16* cKO hearts (N = 4–6 per group). Data are mean ± SD; *p*-values are indicated in each panel. Statistical significance was determined using two-way ANOVA followed by Tukey’s post hoc test.

### *Perm1* gene delivery extends the survival of *Prdm16* cKO mice

We previously showed that patients with *PRDM16* deletion have an increased risk of death, and mice with cardiomyocyte-specific deletion of *Prdm16* have reduced survival^6^. Therefore, we assessed survival up to 40 weeks of age. All AAV9-*Perm1* treated *Prdm16* cKO female mice survived to 20 weeks, whereas only ∼60% of AAV9–GFP treated *Prdm16* cKO female mice were alive at this time point (**Figure 3G**). Up to 40 weeks, AAV9–*Perm1* increased median survival by ∼10–12 weeks; however, mortality emerged after 32 weeks, indicating partial rather than complete long-term rescue (**Figure 3G**). In a separate neonatal cohort, cardiomyocyte-restricted *Perm1* overexpression, driven by the α-myosin heavy chain (αMHC) promoter, similarly improved systolic function in *Prdm16* cKO mice, with significant increases in EF and FS observed 16 weeks post AAV9 delivery **(Figures S2G and S2H**). These findings demonstrate for the first time that *Perm1* gene delivery is protective in the context of genetic cardiomyopathy caused by PRDM16 loss.

### *Perm1* Gene delivery does not restore FA oxidation

Having established that *Perm1* gene delivery improves cardiac function in *Prdm16* cKO mice, we next examined whether it restored FA oxidation. Western blot analysis showed persistent downregulation of HADH and Carnitine palmitoyl transferase 1B (CPT1B) proteins in AAV9-*Perm1*-treated *Prdm16* cKO hearts in both females (**Figures 3H and 3I)** and males **(Figures S3A and S3B**). Consistent with these results, PLC-supported mitochondrial respiration was still reduced in *Prdm16* cKO hearts and was not restored by *Perm1* overexpression in either females (**Figure 3J**) or males (**Figure S3C**). Pyruvate-supported respiration remained unchanged across all groups (**Figure S3D**). These results imply that the cardioprotective effects of *Perm1* are mediated by FAO-independent mechanisms and that PRDM16’s regulation of FAO in the heart involves other regulators.

### PGC1α is required for PRDM16’s regulation of FAO in cardiac cells

Because PERM1 failed to restore FA oxidation *in vivo* despite improving cardiac function, we next asked whether PRDM16 regulates FAO through *Perm1*-independent mechanisms or requires additional transcriptional partners. Given that PGC1α was the only other transcriptional regulator significantly reduced upon PRDM16 loss, we investigated its role in the FAO. Using NRVMs, we showed that Perm1 overexpression increased FA transporter protein expression under basal conditions, but this effect was attenuated when *Prdm16* was depleted (**Figures 4A- 4C**). In contrast, Pgc1a overexpression increased *Perm1* mRNA levels independent of Prdm16 **(Figure 4D)**. Overexpression of Pgc1a restored FAO protein expression even when *Prdm16* was knocked down, indicating that FAO requires Pgc1α (**Figures 4E and 4F**).

**Figure 4.**
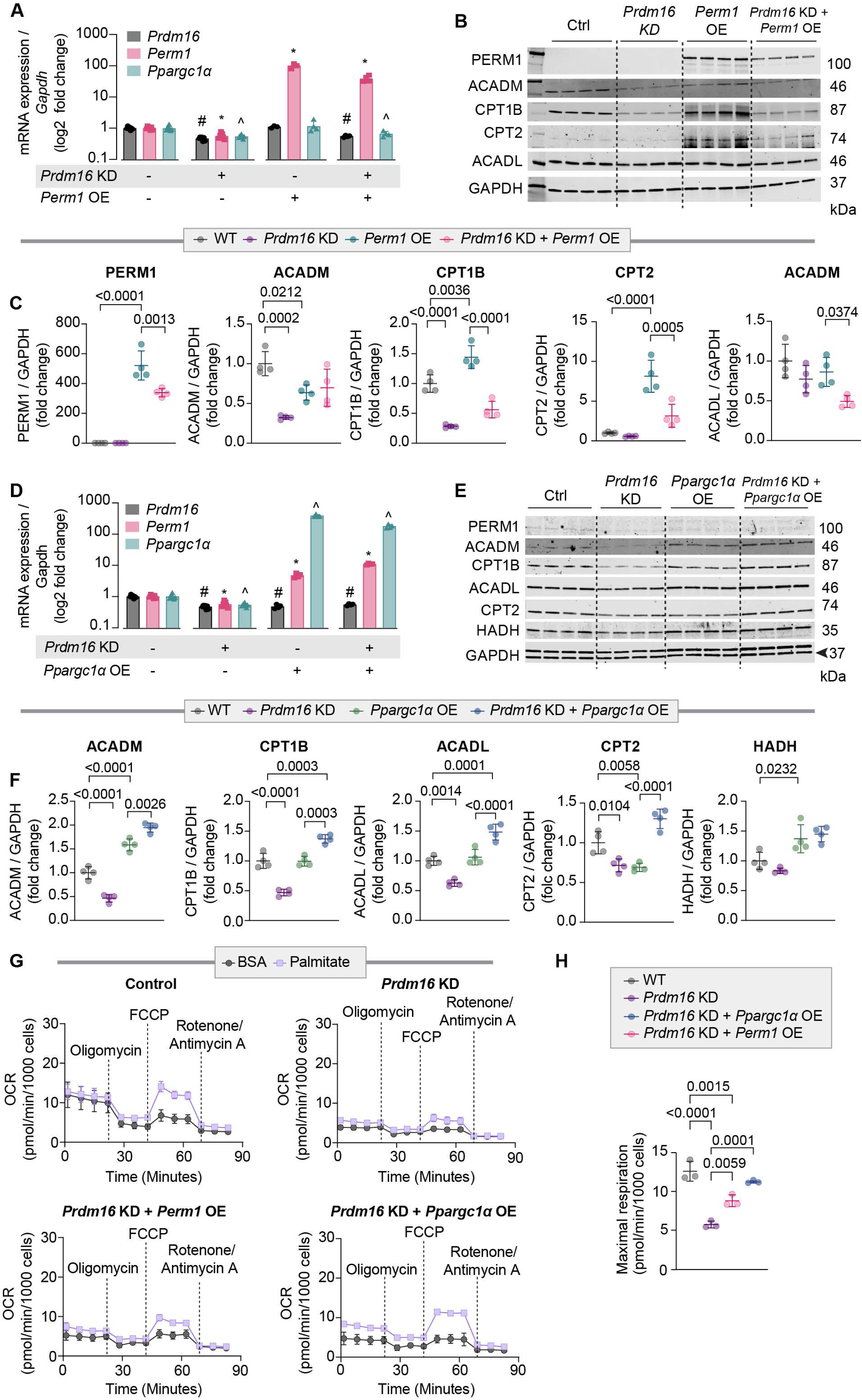
*Perm1* does not restore fatty acid oxidation in cardiac cells in the absence of Pgc1α. (**A**) mRNA expression of *Prdm16*, *Perm1*, and *Ppargc1a* following *Prdm16* knockdown (KD), Perm1 overexpression (OE), or combined *Prdm16* KD + Perm1 OE in NRVMs. Perm1 overexpression markedly increases *Perm1* mRNA expression without altering *Prdm16* or *Ppargc1a* mRNA expression. (**B**) Representative western blots showing protein expression of Perm1 and FAO enzymes under control, *Prdm16* KD, *Perm1* OE, and combined *Prdm16* KD + *Perm1* OE respectively. (**C**) Quantification of protein expression normalized to GAPDH demonstrates that Perm1 overexpression does not fully restore FAO enzyme expression in the setting of *Prdm16* deficiency, despite robust induction of Perm1 (N= 4 per group). (**D**) Relative mRNA expression of *Prdm16*, *Perm1*, and *Ppargc1a* following Prdm16 KD, *Ppargc1a* OE, or combined Prdm16 KD + *Ppargc1a* OE. *Ppargc1a* overexpression significantly increases both *Ppargc1a* and *Perm1* expression, indicating that Pgc1α regulates *Perm1*. (**E-F**) Quantification of FAO-related proteins normalized to GAPDH demonstrates that *Ppargc1a* overexpression restores FAO enzyme protein expression even in the absence of Prdm16 (N= 4 per group). (**G**) Seahorse analysis of mitochondrial palmitate respiration in control and *Prdm16* KD cells, with or without *Perm1* or *Ppargc1a* overexpression. Oxygen consumption rate (OCR) traces show reduced palmitate oxidation following *Prdm16* KD, partial improvement with *Perm1* overexpression, and robust rescue with *Ppargc1a* overexpression. (**H**) Quantification of maximal respiration confirms that Perm1 provides partial improvement, whereas Pgc1α fully restores palmitate oxidation in Prdm16-deficient NRVMs (N= 3 per group). Data is mean ± SD; *p*-values are indicated in each panel. Statistical significance was determined using two-way ANOVA followed by Tukey’s post hoc test.

To directly evaluate the functional consequence of increasing Perm1 and Pgc1a on FAO, NRVMs were transfected with *Prdm16* siRNA or scrambled control and subjected to adenoviral overexpression of Perm1 or Pgc1a prior to assessing palmitate oxidation using the Seahorse XF96 analyzer. Under fatty acid–dependent respiration, overexpression of Pgc1a restored maximal oxygen consumption rate (OCR) to control levels, whereas Perm1 overexpression partially increased it (**Figures 4G and 4H**). These findings indicate that fatty acid–supported mitochondrial respiration requires Pgc1α and cannot be achieved by Perm1 overexpression alone, positioning Pgc1α as a key FAO regulator upstream of Perm1 and downstream of Prdm16 in cardiac cells.

### PERM1 protects from PRDM16-associated cardiomyopathy without affecting the transcriptome

Given that PERM1 improved cardiac function and extended the survival of *Prdm16* cKO mice without restoring FAO, we next examined the mechanisms mediating this protection. RNA sequencing was performed across eight experimental groups, including male and female WT and *Prdm16* cKO mice treated with either AAV9–GFP or AAV9–*Perm1*, respectively. Principal component analysis (PCA) showed clear separation by sex and genotype, with minimal distinction between treatment groups (AAV9–*Perm1* vs. AAV9–GFP) (**Figures S4A- S4B**), indicating limited global transcriptional differences following AAV gene delivery. Volcano plots comparing *Prdm16* cKO hearts treated with AAV9-*Perm1* versus AAV9-GFP in females (**Figure S4C- S4E**) and males (**Figure S4F- S4H)**.

### PERM1 induces post-transcriptional remodeling of mitochondrial and contractile proteomes

The limited transcriptional changes observed following AAV9–*Perm1* treatment, despite robust improvements in cardiac function, suggested that PERM1 may exert its cardioprotective effects predominantly through post-transcriptional mechanisms. We therefore hypothesized that PERM1 regulates cardiac remodeling at the protein level rather than through large-scale transcriptional reprogramming. To directly test this hypothesis, we performed unbiased global proteomic profiling on hearts from female WT and *Prdm16* cKO mice treated with AAV9–GFP or AAV9–*Perm1*, respectively **(Figure S7)**. Principal component (PCA) analysis of the proteomic datasets revealed a clear separation between *Prdm16* cKO and WT hearts. Notably, *Perm1*-treated *Prdm16* cKO hearts shifted toward the WT cluster, indicating partial normalization of the proteomic profile following *Perm1* gene delivery (**Figures 5A and S5A**). Gene ontology cellular component (GO-CC) analysis revealed a large subset of proteins significantly downregulated in the AAV9–GFP-treated *Prdm16* cKO hearts relative to WT hearts, many of which were restored toward WT levels following AAV9–*Perm1* delivery (**Figures 5B, 5C, and S5B**). The significant overlap between pathways dysregulated by loss of PRDM16 and those restored by *Perm1* gene delivery (**Figure 5D and S5C)** indicates that PERM1 selectively reverses a defined subset of PRDM16-dependent proteomic alterations rather than inducing indiscriminate protein expression changes. Importantly, this rescue did not include proteostasis, protein quality control, and canonical FAO enzymes, consistent with the persistent defects in FA-supported mitochondrial respiration (**Figure S5D**). Cluster analysis revealed distinct protein modules that were dysregulated in AAV9–GFP-treated *Prdm16* cKO hearts and selectively restored toward WT levels following *Perm1* overexpression (**Figure 5E**).

**Figure 5.**
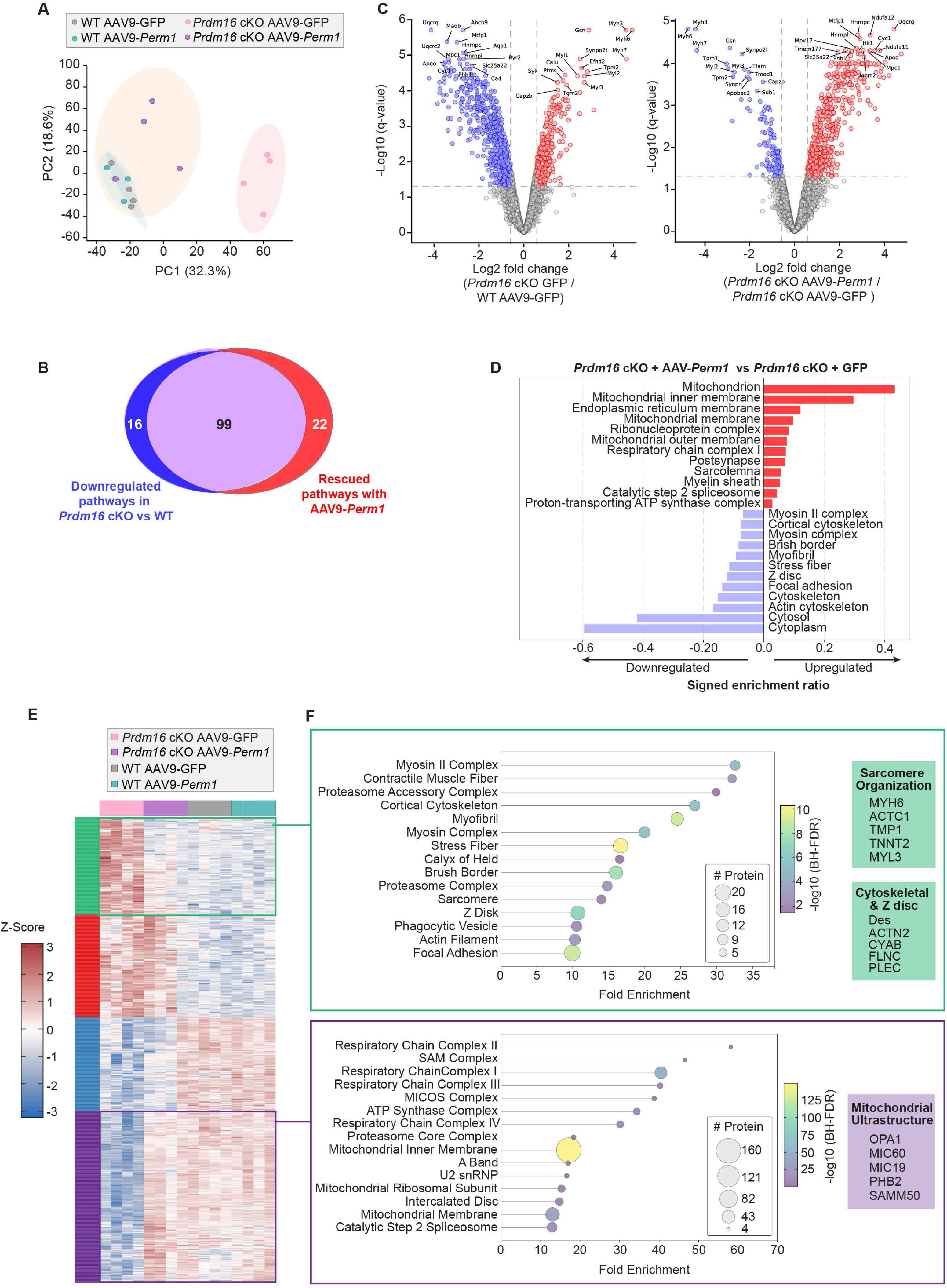
*Perm1* gene therapy induces post-transcriptional remodeling of mitochondrial and cytoskeletal protein networks in *Prdm16* cKO hearts. (**A**) Principal component analysis (PCA) of global proteomic profiles from WT and *Prdm16* cKO hearts from female mice treated with AAV9–GFP or AAV9–*Perm1*. AAV9-*Perm1*-treated *Prdm16* cKO mice shift toward the WT cluster, indicating partial normalization of the proteome. (**B**) Venn diagram showing the overlap between GO Cellular Component (GO-CC) pathways downregulated in *Prdm16* cKO versus WT hearts and those restored by AAV9–*Perm1* gene delivery. (**C**) Volcano plots of differential protein expression highlighting proteins reduced with Prdm16 loss and restored by Perm1 overexpression. (**D**) GO-CC enrichment analysis of proteins altered by Perm1 overexpression in *Prdm16* cKO hearts, ranked by signed enrichment ratio. This analysis identifies upregulation of mitochondrial and membrane-associated compartments and downregulation of cytosolic and cytoskeletal components. (**E**) Unsupervised hierarchical clustering heatmap of differentially expressed proteins across experimental groups reveals distinct protein modules dysregulated in *Prdm16* cKO hearts and partially restored toward WT levels following Perm1 overexpression. (**F**) GO-CC enrichment analysis of *Perm1*-rescued clusters identifies significant enrichment for sarcomere organization, cytoskeletal and Z-disc structures, and mitochondrial structural proteins. Representative proteins include sarcomeric components (MYH6, TNNT2, MYL3), cytoskeletal/Z-disc proteins (ACTN2, FLNC, PLEC), and mitochondrial structural regulators (OPA1, MIC19, PHB2, SAMM50).

GO-CC pathway analysis of the rescued clusters identified significant enrichment for sarcomere organization, cytoskeletal and Z-disc components, and mitochondrial ultrastructural complexes (**Figure 5F**). Sarcomere-associated terms included the myosin II complex, contractile muscle fiber, and myofibril, driven by restoration of key structural proteins such as MYH6, TNNT2, and MYL3. Cytoskeletal and Z-disc–related pathways included cortical cytoskeleton, actin filament, and focal adhesion, with prominent rescue of proteins such as alpha-actinin 2 (ACTN2), filamin C (FLNC), and plectin (PLEC). In parallel, PERM1 overexpression restored multiple components of mitochondrial ultrastructure, including proteins associated with the mitochondrial inner membrane, respiratory chain complexes, and cristae organization. Notably, proteins critical for cristae architecture and mitochondrial integrity, such as optic atrophy 1 (OPA1), mitochondrial contact site and cristae organizing system subunit 19 (MIC19), polyhomeotic homolog 2 (PHC2), and sorting and assembly machinery component 50 homolog (SAMM50), were all significantly enriched within the rescued clusters, suggesting improved mitochondrial structural organization rather than enhanced FA utilization. Together, these data demonstrate that PERM1 overexpression selectively rescues protein networks governing sarcomere integrity, cytoskeletal organization, and mitochondrial ultrastructure. This structural remodeling provides a mechanistic basis for the observed improvement in cardiac function despite a persistent FAO defect.

### *Perm1* gene delivery improves mitochondrial cristae architecture and sarcomere organization in *Prdm16* cKO hearts

To determine whether the *Perm1*-dependent proteomic remodeling translated into ultrastructural improvement, we performed transmission electron microscopy (TEM) on left ventricular tissue from WT and *Prdm16* cKO mice treated with AAV9–GFP or AAV9–*Perm1* (**Figure 6A**). Quantitative analysis demonstrated that mitochondrial size and mitochondrial number were not significantly different among experimental groups (**Figures 6B and 6C**), indicating that PRDM16 loss and PERM1 overexpression do not affect mitochondrial abundance or biogenesis. In contrast, *Prdm16* cKO hearts exhibited a marked reduction in mitochondrial cristae density and irregular cristae organization compared with WT controls (**Figures 6D and 6E**). Importantly, *Perm1* gene delivery significantly increased cristae density and induced a more organized lamellar cristae structure toward WT levels (**Figures 6D and 6E**). In parallel, TEM analysis revealed pronounced sarcomeric disorganization in *Prdm16* cKO hearts, including perturbed sarcomere distribution and Z-disc thickness **(Figure 6F**). *Perm1* overexpression markedly improved sarcomere organization, as evidenced by normalization of sarcomere length distribution (**Figures 6G and 6H**) and improved Z-disc alignment and thickness (**Figure 6I**). Overall, *Perm1* gene delivery improves mitochondrial cristae architecture and sarcomere organization, providing a structural basis for improved cardiac function in *Prdm16* cKO hearts despite persistent impairment in FAO.

**Figure 6.**
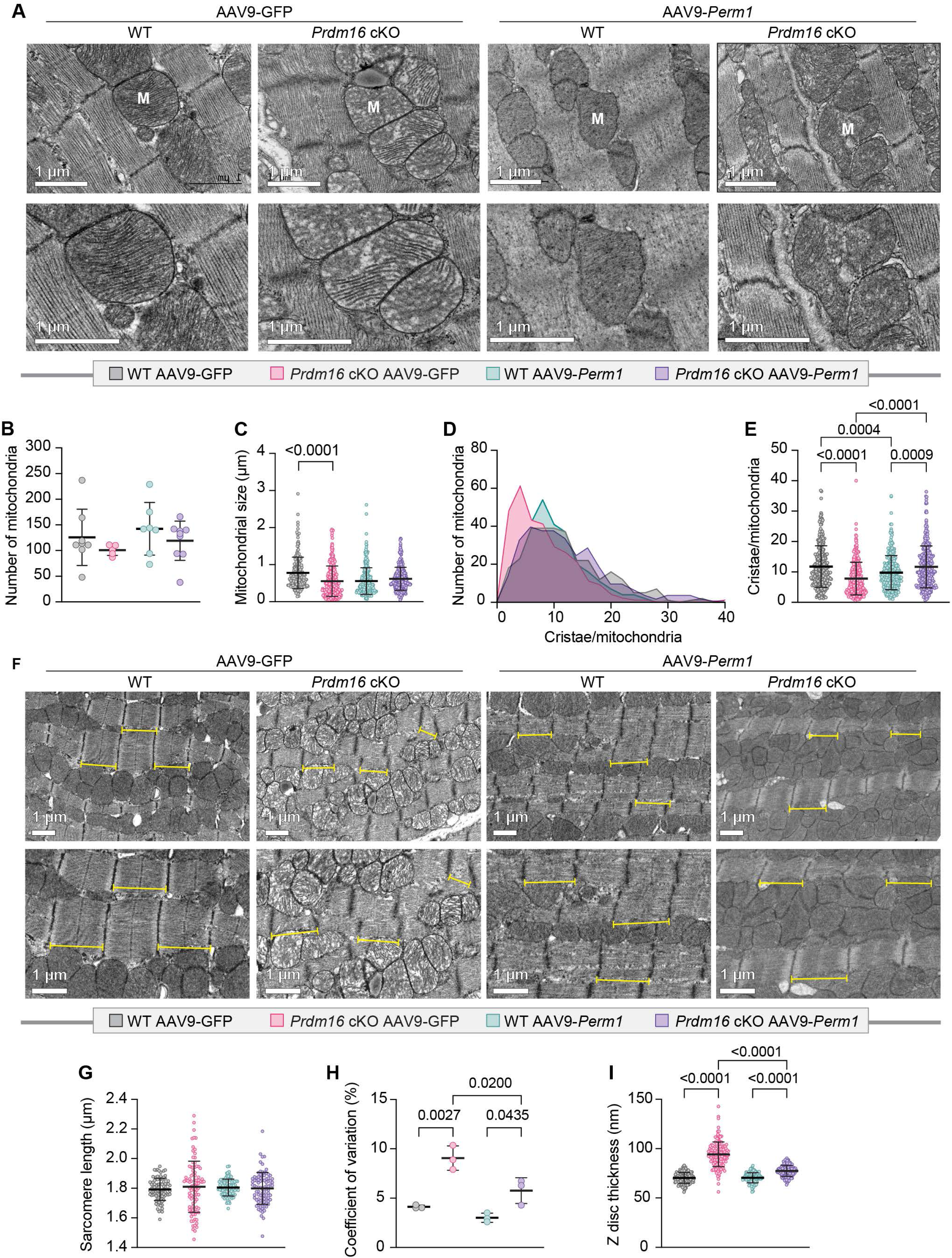
Perm1 improves mitochondrial structure and sarcomere organization in *Prdm16* cKO hearts. (**A**) Representative high-magnification transmission electron microscopy (TEM) images of cardiac mitochondria from hearts of WT and *Prdm16* cKO mice treated with AAV9–GFP or AAV9–*Perm1*. Mitochondria display disrupted cristae architecture in *Prdm16* cKO hearts, which is improved following Perm1 overexpression (scale bars, 1 μm). (**B-C**) Quantification of mitochondrial number and size per field shows no significant differences across experimental groups (N=4 per group). (**D-E**) Distribution and quantification of cristae number (a total of 4 mitochondria per section were quantified) per mitochondrion demonstrate reduced cristae density in *Prdm16* cKO hearts, with a shift toward normalization following *Perm1* gene delivery (N=4 per group). (**F**) Representative low-magnification TEM images illustrating sarcomere organization. *Prdm16* cKO hearts exhibit disrupted myofibrillar alignment and sarcomere structure, which is improved with *Perm1* AAV (scale bars, 1 μm). (**G**) Quantification of sarcomere length shows no major differences in mean sarcomere length across groups. (**H**) Coefficient of variation of sarcomere length reveals increased heterogeneity in *Prdm16* cKO hearts that is significantly improved by Perm1 overexpression. (**I**) Quantification of Z-disc thickness demonstrates increased thickness in *Prdm16* cKO hearts, which is significantly improved with *Perm1* gene delivery. Data is mean ± SD; *p*-values are indicated in the panels. Each data point represents an individual mitochondrion or sarcomere measured from multiple fields per heart across biological replicates. Statistical significance was determined using two-way ANOVA followed by Tukey’s multiple-comparisons test.

### *Perm1* gene delivery reduces cardiomyocyte death, myocardial fibrosis, and cardiomyocyte hypertrophy

Given the improvements in mitochondrial architecture and sarcomere organization following *Perm1* overexpression, we next examined whether these molecular and ultrastructural changes translated into reduced cardiomyocyte death and adverse myocardial remodeling. Cardiomyocyte apoptosis, assessed by TUNEL staining, showed significantly fewer apoptotic nuclei in AAV9–*Perm1*-treated female *Prdm16* cKO hearts compared with AAV9–GFP-treated controls (**Figure 7A**). Consistent with this reduction in apoptosis, western blot analysis of cleaved caspase-3 demonstrated increased cell death in AAV9–GFP-treated *Prdm16* cKO relative to AAV9–GFP-treated WT hearts, which was markedly improved by AAV–*Perm1* delivery **(Figure 7B)**. Given the reduction in apoptotic cell death observed with *Perm1* overexpression, we next evaluated whether cell death prevention was accompanied by alterations in replacement fibrosis. Histological analyses demonstrated a significant decrease in interstitial fibrosis in AAV9–*Perm1*-treated female *Prdm16* cKO hearts as measured by both Picrosirius Red (PSR) and Masson’s Trichrome staining (**Figures 7C and 7D**). Because cardiomyocyte loss and fibrosis are frequently associated with compensatory hypertrophic remodeling in the heart, we quantified cardiomyocyte size. Wheat germ agglutinin (WGA) staining showed a significant reduction in cardiomyocyte cross-sectional area in female *Prdm16* cKO mice treated with AAV9–*Perm1* (**Figure 7E**). These structural improvements were not observed in male mice receiving AAV9–*Perm1* (**Figure S6A-C**). Together, these data demonstrate that *Perm1* gene delivery protects female *Prdm16* cKO hearts from apoptosis, fibrosis, and hypertrophy.

**Figure 7.**
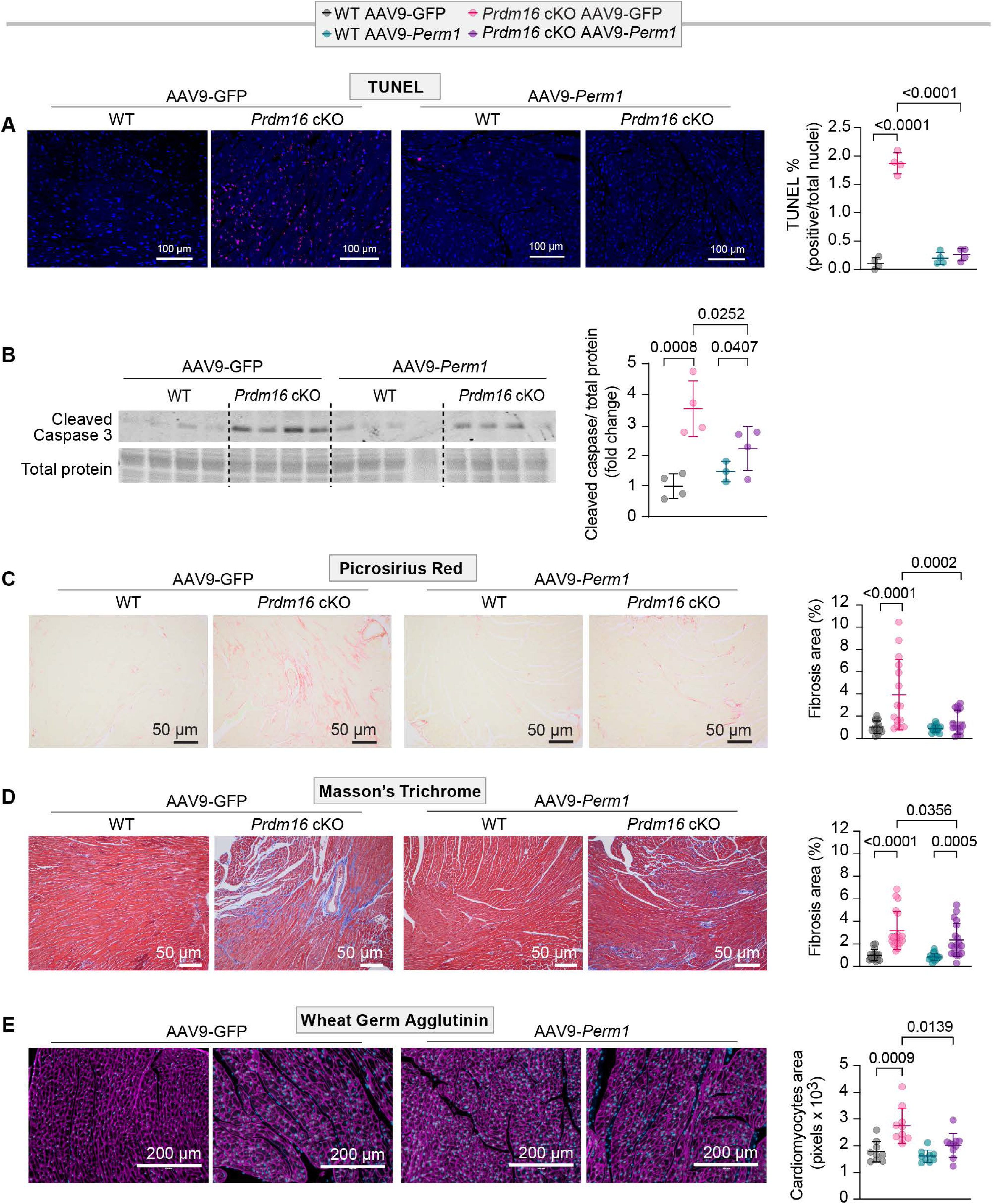
*Perm1* gene delivery attenuates cardiomyocyte death, fibrosis, and cardiomyocyte hypertrophy in female *Prdm16* cKO hearts. (**A**) Representative TUNEL staining of cardiac sections from WT and *Prdm16* cKO mice treated with AAV9–GFP or AAV9–*Perm1*. Quantification shows increased apoptosis in *Prdm16* cKO hearts that is significantly reduced following Perm1 overexpression (scale bars, 100 µm; N = 4 per group). (**B**) Western blot analysis of cleaved Caspase-3 in whole hearts of WT and *Prdm16* cKO mice treated with AAV9–GFP or AAV9–*Perm1*. Quantification normalized to total protein demonstrates elevated apoptotic cell death in *Prdm16* cKO hearts, which is reduced with *Perm1* gene delivery. (**C-D**) Histological assessment of fibrosis by Picrosirius red and Masson’s trichrome staining shows increased collagen deposition in *Prdm16* cKO hearts that is attenuated following AAV9–*Perm1* gene delivery (scale bars, 50 µm; N = 4 per group). (**E**) Wheat germ agglutinin (WGA) staining of cardiomyocytes demonstrates increased cell size in *Prdm16* cKO hearts, with significant reduction following Perm1 overexpression (scale bars, 200 µm; N = 4 per group). Each data point represents measurements from multiple fields per heart across biological replicates. Data is mean ± SD; *p*-values are indicated in the panels. Statistical significance was determined using two-way ANOVA followed by Tukey’s multiple-comparisons test.

## Discussion

Our group and others revealed a previously unrecognized role for PRDM16 in the pathogenesis of LVNC and DCM in humans and mice^6^. In this study, we sought to further understand the mechanisms underlying metabolic dysregulation associated with cardiac PRDM16 deficiency and identify therapeutic targets for PRDM16-associated cardiomyopathy. We showed that cardiomyocyte deletion of *Prdm16* in mice results in an early defect in transcriptional pathways governing FA and oxidative metabolism prior to the development of contractile dysfunction. We identified PERM1 as a downstream transcriptional target for PRDM16 in human and mouse cardiac tissue and cells. We demonstrated a transcriptional regulation of PERM1 by PRDM16, along with the potential direct binding of PRDM16 to the PERM1 promoter region in cardiac cells. These findings establish PERM1 as a downstream target of PRDM16 and a potential therapeutic target.

We observed that PERM1 overexpression improved cardiac function independent of restoring FAO. This was further supported by *in vitro* studies in NRVMs, which revealed that activation of FA metabolic gene expression in the setting of PRDM16 deficiency required PGC1α. Finally, we provide evidence that *Perm1* gene therapy ameliorates PRDM16-associated cardiomyopathy in mice, mainly through post-transcriptional mechanisms involving the preservation of mitochondrial and sarcomere architecture.

PRDM16 belongs to the PR domain family of transcription factors that regulate transcription through intrinsic chromatin-modifying activity and/or by interacting with histone-modifying or other nuclear proteins^24^. In the mouse heart, PRDM16 is expressed in ventricular cardiomyocytes as early as embryonic day 10.5 (E10.5), and its expression is restricted to the compact myocardium^7,9,25^. Consistent with this expression pattern, we and others showed that cardiomyocyte deletion of *Prdm16* in mice caused LVNC and a similar phenotype was observed in patients carrying *PRDM16* deletion/variants^5,6,9,11^. However, we and others also showed that *PRDM16* deletion/variants in humans and mice caused DCM or progressive HF phenotypes^5,7-11,26^, suggesting that this transcription factor regulates genes involved in maintaining energy supply and sarcomere organization.

One of the earliest defects we observed in *Prdm16* cKO male and female mice, prior to the development of overt contractility defects, is a decrease in FAO. While this defect was previously observed in transcriptional and metabolomic profiling in other *Prdm16* cKO mice^7-11^, the mechanisms driving this decrease in FAO were not thoroughly investigated. This is the first study to show that FAO gene network is regulated perinatally by PRDM16 in the mouse heart. In fact, we investigated several transcriptional regulators of FA metabolism in the heart, including *Pparα*, *Ppargc1a*, *Errα/Errγ*, and *Perm1*^19,27-29^, and found that only *PPERM1* and *PPARGC1A* expression was consistently decreased with PRDM16 deficiency in both iPSC-CMs from a proband carrying the *PRDM16Q187X* variant and the hearts of *Prdm16Q187X* knock-in (KI) mice. These findings position PGC1α and PERM1 as direct downstream effectors of PRDM16-dependent FAO metabolism.

Across multiple experimental systems, we showed that PRDM16 deficiency consistently reduced PERM1 expression. Unlike PGC1α, which controls broad metabolic programs and is often regulated by stress, nutrient status, and developmental cues, PERM1 provides a more direct mechanistic link to FA metabolism, as it has been shown to directly bind *Pparα* and drive FAO gene expression specifically in the mouse heart^21^. Additionally, Perm1 is exclusively expressed in skeletal and heart muscle and to a lesser degree in BAT in mice. Perm1 uniquely integrates transcriptional and structural control of mitochondrial bioenergetics, including interactions with the mitochondrial contact site and cristae organizing system–mitochondrial intermembrane space bridging (MICOS–MIB) complex that are not shared by PGC1*α* or PPARα^20^. In addition, sustained Pgc1*α* overexpression in the mouse heart induced maladaptive remodeling, limiting its therapeutic utility^15-17^. Based on these features, we postulated that Perm1 may constitute an ideal candidate for restoring cardiac FAO deficits observed in *Prdm16* cKO mice. AAV9-mediated *Perm1* overexpression produced a remarkable functional improvement, evidenced by increased EF and FS, limited ventricular dilation, and significantly enhanced long-term survival in *Prdm16* cKO female mice. Importantly, this benefit was observed 20-40 weeks after a single neonatal injection of AAV9–*Perm1*, highlighting long-term biological effects. Compared to recent work showing that *Perm1* overexpression improved cardiac function in healthy adult mice^30^ and pressure overload HF mouse models^22,23^, we extended these findings by demonstrating the long-term therapeutic benefit of *Perm1* gene therapy in Prdm16-associated cardiomyopathy in mice.

A particularly unexpected finding in our study was that *Perm1* improved cardiac performance without restoring FAO capacity. Cardiac expression of FAO enzymes remained suppressed, and mitochondrial oxidation of PLC was still impaired in AAV9–*Perm1*-treated mice. We hypothesized that PRDM16 regulates FA metabolic gene expression through the PGC1α–PERM1 axis. This model is supported by our *in vitro* studies on NRVMs showing that *Perm1* overexpression alone partially increased protein expression of a subset of FA enzymes and elevated palmitate oxidation, whereas Pgc1*α* overexpression completely rescued the FAO defect in PRDM16-deficient cardiac cells. These findings indicate that although PERM1 contributes to FA metabolic regulation, it is not sufficient to rescue the FAO defect in the absence of PGC1α.

Our RNA-seq analysis revealed limited transcriptional changes following Perm1 overexpression. This indicates that PERM1 functional benefits do not arise from restoration of canonical FAO transcriptional pathways. This dissociation between transcriptional remodeling and functional rescue suggests that PERM1-mediated cardioprotection operates at the post-translational level. Indeed, unbiased global proteomic profiling revealed that *Perm1* overexpression selectively rescued discrete protein networks disrupted by PRDM16 loss, rather than globally normalizing the cardiac proteome. These rescued networks were enriched for proteins governing sarcomere organization, cytoskeletal and Z-disc integrity, and mitochondrial ultrastructure, whereas canonical FAO enzymes remained largely downregulated. Importantly, this interpretation aligns with emerging literature showing that PERM1 interacts with cytoskeletal and mitochondrial-associated proteins, supports sarcomere organization, stabilizes mitochondrial microdomains, and modulates calcium handling^20,31^. This selective proteomic remodeling supports a model in which PERM1 stabilizes structural and organellar protein assemblies that buffer cardiomyocytes against stress without restoring metabolic capacity. Importantly, these proteomic signatures translated directly into ultrastructural improvements at the organelle and myofibrillar levels, as demonstrated by TEM. Although mitochondrial number and size were unchanged across groups, *Prdm16* cKO hearts exhibited marked disruption of cristae density and organization, indicative of compromised mitochondrial inner membrane architecture. *Perm1* gene delivery significantly restored cristae density and lamellar organization toward WT levels, consistent with proteomic enrichment of cristae-associated and MICOS–MIB complex proteins. In parallel, TEM analysis revealed pronounced sarcomere disorganization in PRDM16-deficient hearts, including irregular sarcomere alignment and abnormal Z-disc morphology. Perm1 overexpression normalized sarcomere length distribution and improved Z-disc alignment and thickness, providing ultrastructural confirmation of the rescued sarcomere and cytoskeletal protein networks identified by proteomics.

Together, these RNA-seq, proteomic, and ultrastructural data define a mechanism by which PERM1 suppresses stress signaling and preserves cardiomyocyte integrity by stabilizing mitochondrial and sarcomere architecture rather than recovering FA metabolism, which, in the case of PRDM16 deficiency, may need a functional PGC1α. By improving mitochondrial ultrastructure and force-transmitting cytoskeletal networks, PERM1 likely reduces mechanical stress that would otherwise trigger DNA damage responses, p53 activation, and maladaptive cell-cycle re-entry. Support for this model came from complementary assays demonstrating that PERM1 substantially attenuates cardiomyocyte death and adverse remodeling. These findings are consistent with prior work showing that Perm1 overexpression limited hypoxia/reoxygenation-induced apoptosis by preserving mitochondrial stability and reducing oxidative stress^19^. PERM1 also mitigated pathological remodeling in PRDM16-deficient hearts. Notably, restoration of sarcomere alignment and cytoskeletal anchoring provides a mechanistic basis for this antifibrotic effect, as improved intracellular force distribution is known to limit profibrotic mechano-transduction signaling^32^. These findings are consistent with prior *in vivo* studies showing that Perm1 overexpression reduces myocardial fibrosis in pressure–overload-induced HF and in AAV-mediated therapeutic models, where improved cardiac function is accompanied by decreased extracellular matrix accumulation^33^. Furthermore, recent structural studies have demonstrated that PERM1 interacts with components of the MICOS–MIB complex and ankyrin B to link mitochondria to the cytoskeleton and sarcolemma^20^, supporting a role in maintaining subcellular organization under mechanical stress.

Sex-specific responses to PERM1 represent another important finding of this study. Female *Prdm16* cKO mice exhibited clear and durable functional improvement following Perm1 overexpression, whereas male *Prdm16* cKO mice showed neither harm nor significant improvement, which we postulate is due, in part, to the detrimental effect of AAV9–Perm1 in male WT mice. Indeed, long-term Perm1 overexpression was associated with a 5-10% reduction in EF in male WT mice, suggesting that biological sex may influence the cardiac responses to sustained Perm1 overexpression. Alternative contributors to sex-specific effects of AAV9–*Perm1* may include sex differences in mitochondrial signaling and stress responses, technical variables such as sex-dependent variation in AAV transduction efficiency, and sex differences in immune responses^34-36^. Further studies are needed to determine the underlying mechanisms of sex differences in cardioprotection mediated by *Perm1* gene therapy.

This study has limitations. Our study relies on neonatal AAV9 delivery, which yields robust and long-lasting cardiac transduction but may not fully reflect therapeutic scenarios in juvenile or adult-onset cardiomyopathy. Neonatal administration can produce biological responses distinct from those activated when gene therapy is delivered after myocardial remodeling has already begun. Whether Perm1 overexpression initiated later in life would confer similar benefits remains to be determined.

In conclusion, this study identifies a previously unrecognized PRDM16–PGC1α–PERM1 signaling axis that is essential for maintaining FAO in the heart. *Perm1* gene therapy significantly improves cardiac performance and survival by reducing cell death, attenuating fibrosis, and preventing cardiomyocyte hypertrophy in *Prdm16* cKO female mice. These findings demonstrate that PERM1 confers cardioprotection through mechanisms independent of FAO and highlights the structural stabilizing roles for PERM1 in the heart. Collectively, our results position PERM1 as a promising therapeutic modifier for PRDM16-associated cardiomyopathy. Future studies will explore whether *PERM1* gene therapy will ameliorate mitochondrial and sarcomere organization in iPSC-CMs and engineered heart tissue from patients carrying deleterious *PRDM16* variants.

## Non-standard Abbreviations and Acronyms

AAV: Adeno-associated virus
cKO: Cardiac knockout
ChIP: Chromatin immunoprecipitation
CMs: Cardiomyocytes
FAO: Fatty acid oxidation
FS: Fractional shortening
NRVMs: Neonatal rat ventricular myocytes
OCR: Oxygen consumption rate
P1: Postnatal day 1
PLC: Palmitoyl-L-carnitine
PSR: Picrosirius red
TUNEL: Terminal deoxynucleotidyl transferase dUTP nick-end labeling

## Acknowledgments

We are grateful to our collaborators for their guidance and technical expertise. We thank Ms. Allison Mannuel and the Metabolomics Core Facility at the University of Utah for their assistance with the proteomics analysis. We also thank Ms. Diana Lim and Ms. Nikita Abraham for assistance with figure preparation. We are grateful to Dr. Junco Warren for sharing the the PERM1 plasmid and for providing whole-heart homogenate from cardiomyocyte-specific *Perm1* knockout mice to use as a control in western blot.

---

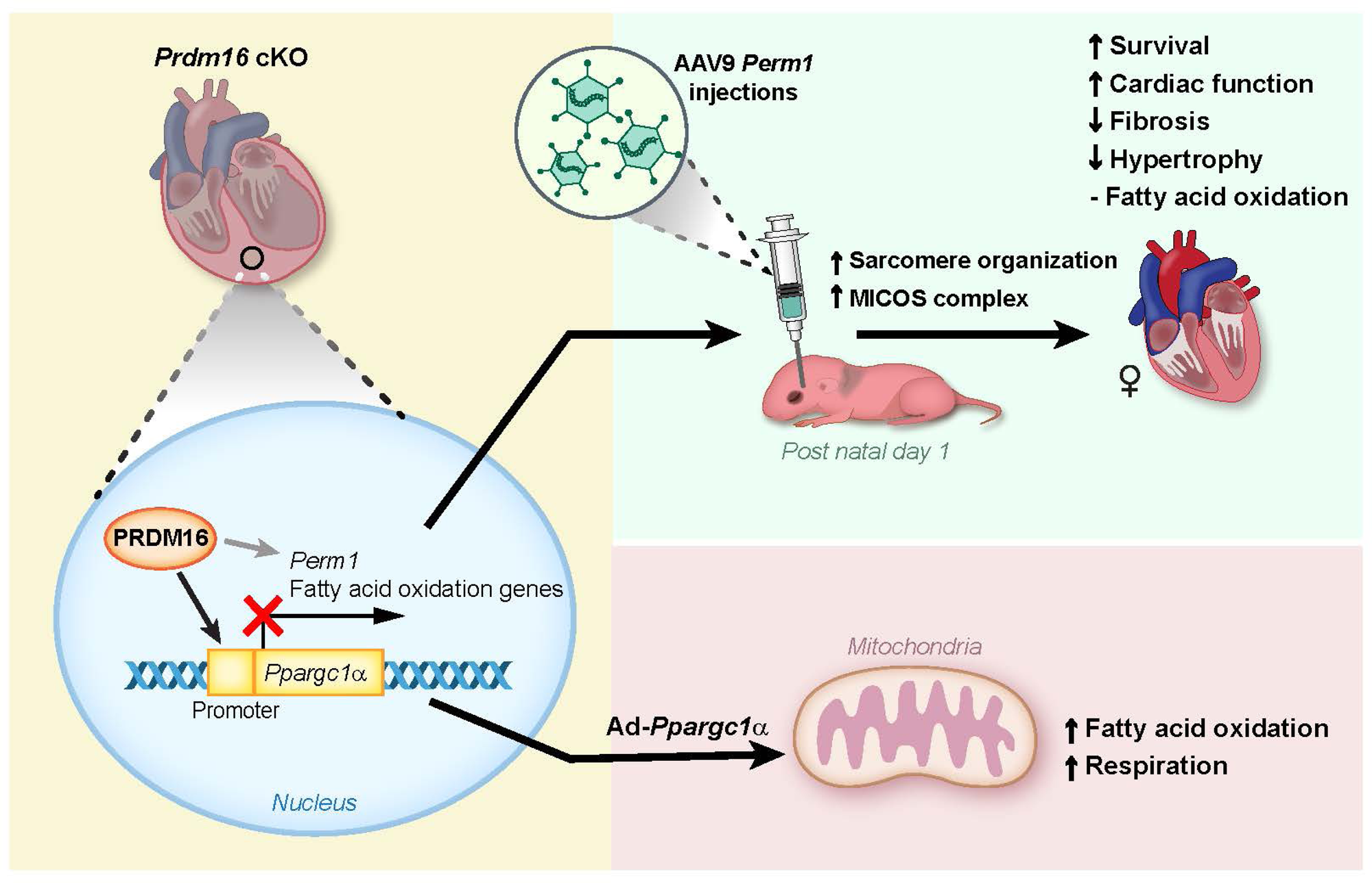

